# Measurement error and variant-calling in deep Illumina sequencing of HIV

**DOI:** 10.1101/276576

**Authors:** Mark Howison, Mia Coetzer, Rami Kantor

## Abstract

**Motivation:** Next-generation deep sequencing of viral genomes, particularly on the Illumina platform, is increasingly applied in HIV research. Yet, there is no standard protocol or method used by the research community to account for measurement errors that arise during sample preparation and sequencing. Correctly calling high and low frequency variants while controlling for erroneous variant calls is an important precursor to downstream interpretation, such as studying the emergence of HIV drug-resistance mutations, which in turn has clinical applications and can improve patient care.

**Results:** We developed a new variant-calling pipeline, hivmmer, for Illumina sequences from HIV viral genomes. First, we validated hivmmer by comparing it to other variant-calling pipelines on real HIV plasmid data sets, which have known sequences. We found that hivmmer achieves a lower rate of erroneous variant calls, and that all methods agree on the frequency of correctly called variants. Next, we compared the methods on an HIV plasmid data set that was sequenced using an amplicon-tagging protocol called Primer ID, which is designed to reduce errors and amplification bias during library preparation. We show that the Primer ID consensus does indeed have fewer erroneous variant calls compared to the variant-calling pipelines, and that hivmmer more closely approaches this low error rate compared to the other pipelines. Surprisingly, the frequency estimates from the Primer ID consensus do not differ significantly from those of the variant-calling pipelines. Finally, we built a predictive model for classifying errors in the hivmmer alignment, and show that it achieves high accuracy for identifying erroneous variant calls.

**Availability:** hivmmer is freely available for non-commercial use from https://github.com/mhowison/hivmmer.

## 1 Introduction

Several next-generation sequencing instruments are now used to study pathogens and viruses (Chabria et al., 2014; Quiñones-Mateu *et al.*, 2014). Of the many next-generation sequencing platforms and approaches that have been developed over the past two decades, Illumina’s sequencing-by-synthesis technology has come to dominate the market, in large part due to increasing yields and decreasing costs (Goodwin *et al.*, 2016). Deep sequencing of HIV samples with Illumina technology is frequently used in studies of viral epidemiology, clinical genotyping, and antiretroviral drug resistance. For example, deep sequencing can provide for a more sensitive assay of drug-resistance mutations (Brumme and Poon, 2016); Sanger sequencing, the current clinical standard, cannot reliably detect mutations at frequencies below 20%, which might be clinically relevant (Ávila Ríos *et al.*, 2016). A common concern in studies using deep sequencing, and also in establishing clinical standards for these new approaches, is the measurement error of their sequencing protocols. Measurement errors can arise in sample preparation (including reverse transcription of RNA genomes to cDNA and amplification of viral genomes), library preparation, sequencing and base calling.

Measurement error creates uncertainty in downstream analyses. For example, errors introduced during genome amplification are difficult to distinguish from real mutations since they are introduced in the early steps of the process, are exponentially amplified and may occur at high frequency in the later steps. Recombination during PCR is difficult to distinguish from clinically-relevant “real” viral recombination. Mutations at low frequencies can be difficult to distinguish from sequencing and base calling errors, and can confound read alignment, assembly and haplotype reconstruction methods that rely on accurately identifying exact sequence overlaps among sequence reads. Beerenwinkel *et al.* (2012) speculated that artifacts introduced during the RT-PCR step are likely the biggest challenge to accurately estimating viral diversity through reconstructing individual haplotypes for deeply sequenced HIV data.

Many HIV studies in recent years have addressed Illumina sequencing errors by applying a global frequency threshold – typically 1% – below which variants are excluded with the reasoning that they are indistinguishable from sequencing errors. This approach requires establishing a conservative estimate of the typical error rate for the sequencing protocol, which is then used as a threshold during variant calling.

The most common approach to estimating sequencing error rates is to analyze reads that come from known sequences, by aligning the reads to the known sequence and counting the frequency of mismatches in the alignment. In the context of HIV, this can be accomplished by sequencing mixtures of HIV plasmids with known sequences. In this study, we use this approach to introduce a new pipeline for HIV *pol* sequences from the Illumina MiSeq platform, hivmmer, and compare it to other variant-calling pipelines. While existing pipelines use short-read aligners to align Illumina reads in nucleotide space against an HIV reference (such as HXB2; accession K03455) or a *de novo* assembly, hivmmer instead uses a probabilistic aligner, HMMER (Eddy, 2011), to achieve a more sensitive alignment in amino-acid space.

## 2 Methods and Data

### 2.1 Pipelines

We created a new pipeline, hivmmer (version 0.1.0), based on the probabilistic aligner HMMER (Eddy, 2011), that consists of the following steps:

1. Constructs an amino acid profile Hidden Markov Model (pHMM) from a multiple sequence alignment of all HIV-1 Group M amino acid sequences publicly available in the Los Alamos HIV Sequence Database (http://www.hiv.lanl.gov) for the *pol* gene.
2. Preprocesses the NGS data using the paired-end read merging tool PEAR (Zhang *et al.*, 2014) and consolidates duplicate sequences using FASTX-Toolkit (http://hannonlab.cshl.edu/fastx_toolkit/). The number of duplicates are tracked to enable correct inference of frequencies later in the pipeline.
3. Translates each de-duplicated sequence into all six possible frames (forward and reverse), retaining only the translated sequences that contain no stop codons.
4. Aligns the translated reads to the reference pHMM with hmmsearch from HMMER, producing a multiple sequence alignment of translated reads.
5. Constructs a sample-specific amino acid pHMM from the multiple sequence alignment of translated reads.
6. Repeats the hmmsearch alignment against the sample-specific pHMM for increased sensitivity.
7. Maps the translated amino acid coordinates in the alignment to the original frame and coordinates in the nucleotide reads to construct a codon frequency table (adjusting the counts for duplicate reads).

We compared hivmmer to two of the existing pipelines, HyDRA (Ji *et al.* (2015); version 1.2.6-a41ac1f) and shiver (Wymant *et al.* (2016); version 1.3.0), both of which use the short-read aligner bowtie2 (Langmead and Salzberg, 2012). HyDRA aligns the reads to the HXB2 reference, while shiver uses an iterative alignment to a *de novo* assembly of the reads. We chose HyDRA and shiver as representatives of a broader group of HIV alignment pipelines such as PASeq (https://paseq.org) and MiCall (http://cfe-lab.github.io/MiCall), which are also based on bowtie2 (for a recent comparison of these methods, see Noguera-Julian *et al.* (2017)).

## 2.2 Data

Our study uses four publicly-available HIV plasmid data sets:

1. **5VM** (accession SRR961514), a “5 virus mix” of plasmid sequences (89.6, HXB2, JRCSF, NL4-3, YU2) in equal proportions (20%) sequenced by Di Giallonardo *et al.* (2014);
2. **PL1:1** (accession SRR6725661), a mixture of two plasmid sequences in 1:1 proportion generated in our lab and introduced in this study (described below).
3. **PL1:9** (accession SRR6725662), the same mixture as PL1:1, but in 1:9 proportion;
4. **PID** (accessions SRR2097103-8), the same mixture as 5VM, but sequenced using the Primer ID protocol (discussed in depth below) by Seifert *et al.* (2016).

For PL1:1 and PL1:9, plasmids pNL4.3 (AF324493.2) and p89.6 (U39362), obtained from the NIH AIDS Reagent Program (https://www.aidsreagent.org/), were mixed as 1:1 or 1:9 ratios respectively, followed by amplification of the *pol* region using primers previously described by Winters *et al.* (1998), and proof reading polymerase Phusion (Thermofisher). Nextera XT DNA Library Prep chemistry (Illumina) was used to fragment and add adapter sequences onto template DNA to generate multiplexed sequencing libraries that were sequenced on Illumina’s MiSeq platform generating 2 x 250bp paired-end reads.

5VM contains near-full-length HIV genomes, although for this study we considered their alignment and variant calls only within the first 1044nt of the *pol* region (HXB2 coordinates 2253-3296). This is also the region contained in PL1:1 and PL1:9, and is a genomic region that is clinically relevant for drug resistance mutations. PID is a restricted fragment within this region, with length 471nt starting at HXB2 coordinate 2736.

The sequencing method used to generate PID was initially developed by Jabara *et al.* (2011) to tag each individual HIV viral template with a unique “Primer ID” prior to PCR. Jabara *et al.* (2011) demonstrated with Roche 454 sequencing that this technique can be used to construct consensus sequences that accurately reflect the original input templates without PCR artifacts. Keys *et al.* (2015) extended this approach to Illumina MiSeq sequencing and demonstrated improvements in detecting drug resistance mutations. Zhou *et al.* (2015) conducted additional statistical modeling to refine the Primer ID method with Illumina MiSeq sequencing and address technical issues, such as the effect of sequencing errors in the primer sequence itself. They found that greater than 90% of the sequence reads were useable. In contrast, a study by Brodin *et al.* (2015) with 454 Roche sequencing found that PCR errors in the Primer ID sequences led to skewed resampling and in some cases 99% of the original input templates were not recoverable as consensus sequences. However, a study by Seifert *et al.* (2016) with Illumina MiSeq sequencing of plasmid sequences found that the loss of reads due to incorrect Primer IDs was only 3%.

The PID data set comes from this latter study by Seifert *et al.* (2016). We compared the variants calls in the consensus sequences from their Primer ID aligner, called pidalign, to those from running each of the pipelines on the original reads with the Primer ID barcodes removed. That is, we tested the pipelines under the condition where the Primer ID is unknown.

## 2.3 Machine learning for error classification

Machine learning refers to the use of statistical and algorithmic techniques that learn patterns from a set of training data to make predictions for a new set of previously unseen data (Hastie *et al.*, 2009). A common application of machine learning is classification, meaning to predict to which of a finite set of classes a data point is most likely to belong. In the context of sequencing errors, data points could be individual base calls, alignments, or variant calls; and the classification task can be classifying them as either errors or non-errors.

We trained and validated a classifier for the variant calls in hivmmer, using a gradient boosting method called xgboost (Chen and Guestrin, 2016). Briefly, gradient boosting is a machine learning algorithm that constructs an ensemble of decision trees from training data, and makes predictions on new data by averaging the paths through the decision trees. We constructed the following predictive features for each variant call in the final hivmmer alignment: position of call in the read, read length, read frequency (from the de-duplication count), position of call in the alignment, alignment length, HMMER’s posterior probability for the call, and dummy variables for the aligned frame. We used a cross-validation approach in which we trained a model for each dataset (5VM, PL1:1, PL1:9) and validated it on the opposite type. We also evaluated the model for each dataset on itself, using a randomly sampled 50% hold-out validation set.

## 2.4 Reproducibility

All scripts required to reproduce the results presented here are available from https://github.com/mhowison/hiv-measurement-error and can be executed using an accompanying biomake file (Holmes and Mungall, 2017). Compiled versions of all software dependencies for 64-bit Linux and Anaconda Python (https://www.anaconda.com) are available from the hivmmer conda channel (http://anaconda.org/hivmmer). The hivmmer source code is available from (https://github.com/mhowison/hivmmer) and a pre-compiled Docker (https://docker.com) image is available from DockerHub at https://hub.docker.com/r/mhowison/hivmmer.

## 3 Results

We analyzed the coverage and fragment sizes of the Illumina reads. Figure 1(a) shows an overview of fragment sizes in 5VM; PL1:1 and PL1:9 have similar fragment size distributions. Typically, fragment sizes follow a skewed distribution centered around the read length, 250nt. Fragments shorter than the read length are fully overlapping, and yield reads that in practice can be treated as technical replicates. Fragments sized between the read length and twice the read length yield partially overlapping reads that can be combined into a single sequence using a read-merging tool like PEAR (Zhang *et al.*, 2014). Finally, fragments larger than twice the read length yield separate read pairs, with a positive insert size between the reads. The coverage of the reads across the genome tends to be variable (Fig. 1b), as does the relative proportion of replicate, overlapping, and paired reads. For example, in 5VM there are relatively fewer paired reads observed in highly variable regions such as *env* (HXB2 coordinates 6045-8795).

**Figure 1.**
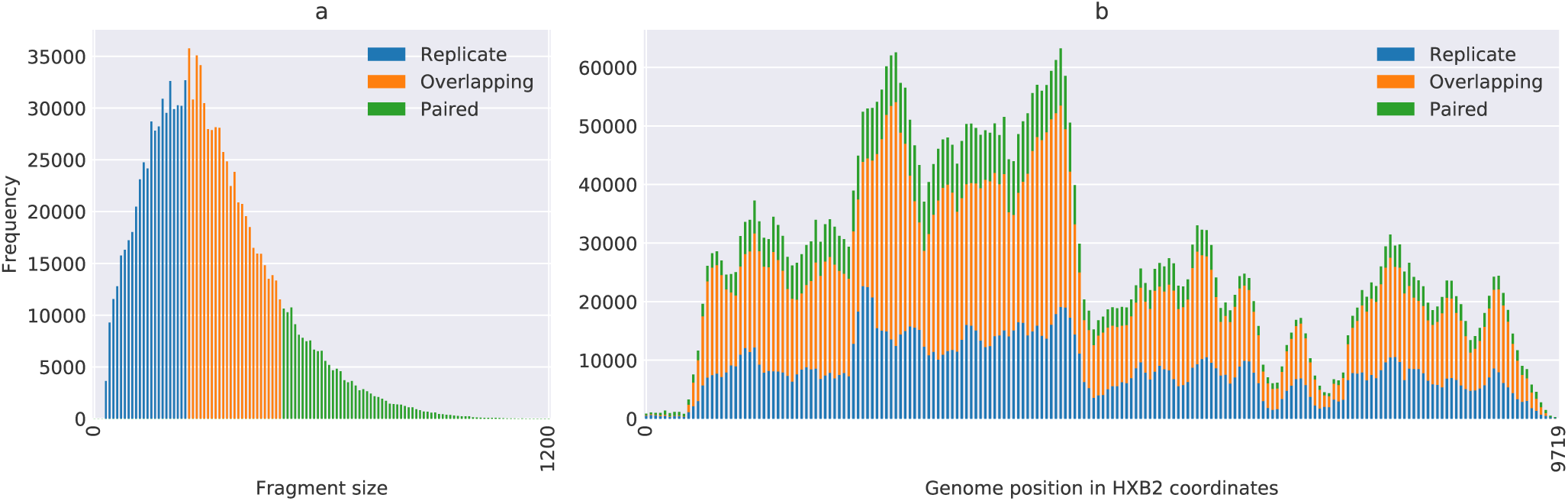
Overview of read distribution frequencies from the 5VM data set. Fragment size frequencies (a) illustrate the proportion of reads that are either technical replicates (fragment is less than the maximum read length; blue), overlapping (fragment is between the read length and twice the read length; orange), or paired-end (fragment is larger than twice the read length; green). Coverage (b) is non-uniform across the HIV genome due to the use of multiple amplicons to cover the whole genome.

To compare pipelines, we first identified both the correct and erroneous calls in the underlying alignments from each pipeline. We defined erroneous calls as codons with ¿0 frequency, but which do not exist at that position in any of the known plasmid sequences for the given data sets. Supplementary Figures 1-4 show a detailed picture of this for each data set and pipeline. As expected, nearly all of the erroneous calls are at frequencies below 1%, which is a widely accepted global threshold. However, a few calls for HyDRA and shiver on 5VM are above 1%. To measure the overall effectiveness of each pipeline, we plotted the cumulative number of errors as we lowered the global frequency threshold from 2% to 0.1% (Figure 2). From this plot, we find that hivmmer alignments accumulate fewer errors across all data sets.

**Figure 2.**
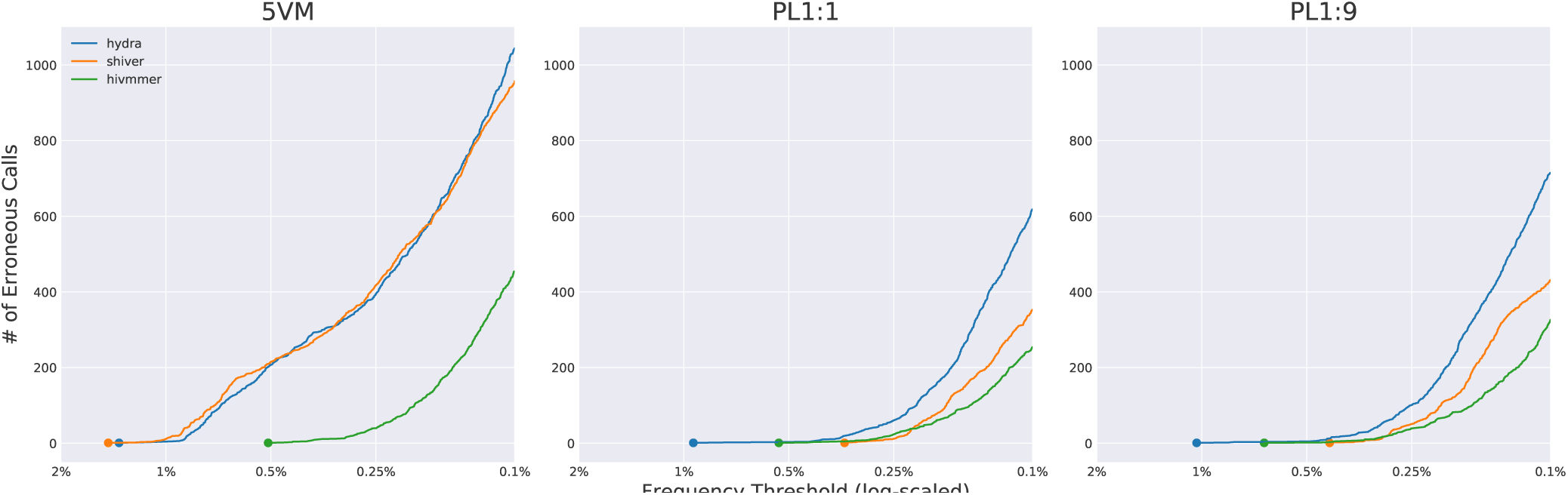
Comparison of cumulative errors observed in the alignments, across data sets and methods. For 5VM, hivmmer alignments display the lowest cumulative error rate. For PL1:1 and PL1:9, the cumulative error rates are closer among the methods, but hivmmer alignments display fewer errors at thresholds below 0.25%.

Next, we considered the frequencies of correctly called variants, and compared their distribution across pipelines (Figure 3). While we expect the frequencies to follow the mixture proportions (e.g. multiples of 20% for 5VM, 1:1 for PL1:1, and 1:9 for PL1:9), in reality the frequencies deviate from these expected values. This could be due to sample preparation or preferential primer amplification. Because we have no ground truth, we cannot assess which pipeline is most accurate. However, we can test the null hypothesis that at least one of the distributions is significantly different from the others using the Kruskal-Wallis test, a non-parametric analog to the ANOVA. This test fails to reject the null for any of the data sets (5VM *p* = 0.891; PL1:1 *p* = 0.996; PL1:9 *p* = 1.000). Visual inspection of Figure 3 confirms this, as the distributions are nearly indistinguishable. Therefore, none of the pipelines differed significantly in their measurement of the variant frequencies for correctly called variants.

**Figure 3.**
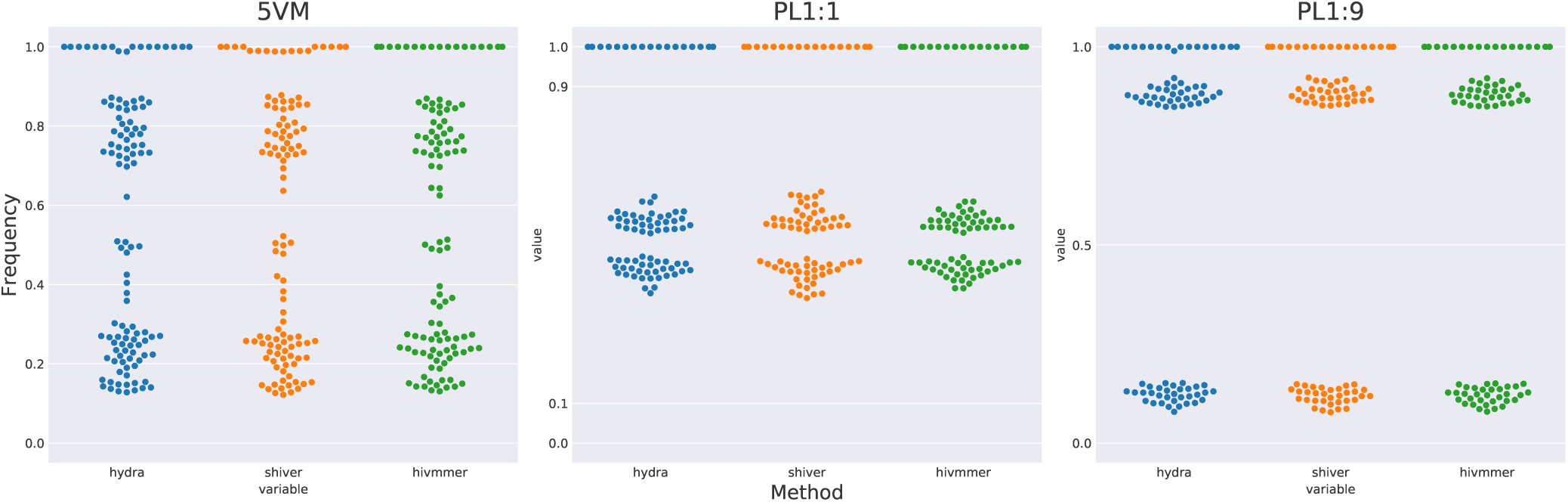
The distribution of variant calls across data sets and methods. Visually, the distributions do not appear significantly different across methods, which is confirmed using a Kruskal-Wallis test.

The Primer ID protocol was designed to control for the artifacts during sample preparation that could be potentially skewing our recovered frequencies of correctly called variants. We compared the cumulative error rate and distribution of call frequencies between two of the pipelines (HyDRA and hivmmer, as the initial *de novo* assembly of the PID data set failed for shiver) and the pidalyse method for calling the consensus sequence of each Primer ID template (Seifert *et al.*, 2016). Because these consensus sequences should represent individual templates, we expect that the frequency of calls across templates would correspond to the plasmid mixture proportions (e.g. multiples of 20% for 5VM). The Primer ID method does indeed reduce the accumulation of erroneous calls, and hivmmer better approaches this performance than HyDRA (Fig. 4a). The Primer ID consensus sequences, however, also do not recover the correct frequency proportions as one might expect (Fig. 4b). This is consistent with the results reported by Seifert *et al.* (2016), and they ascribe the discrepancy to “noisy RT qPCR quantification.”

**Figure 4.**
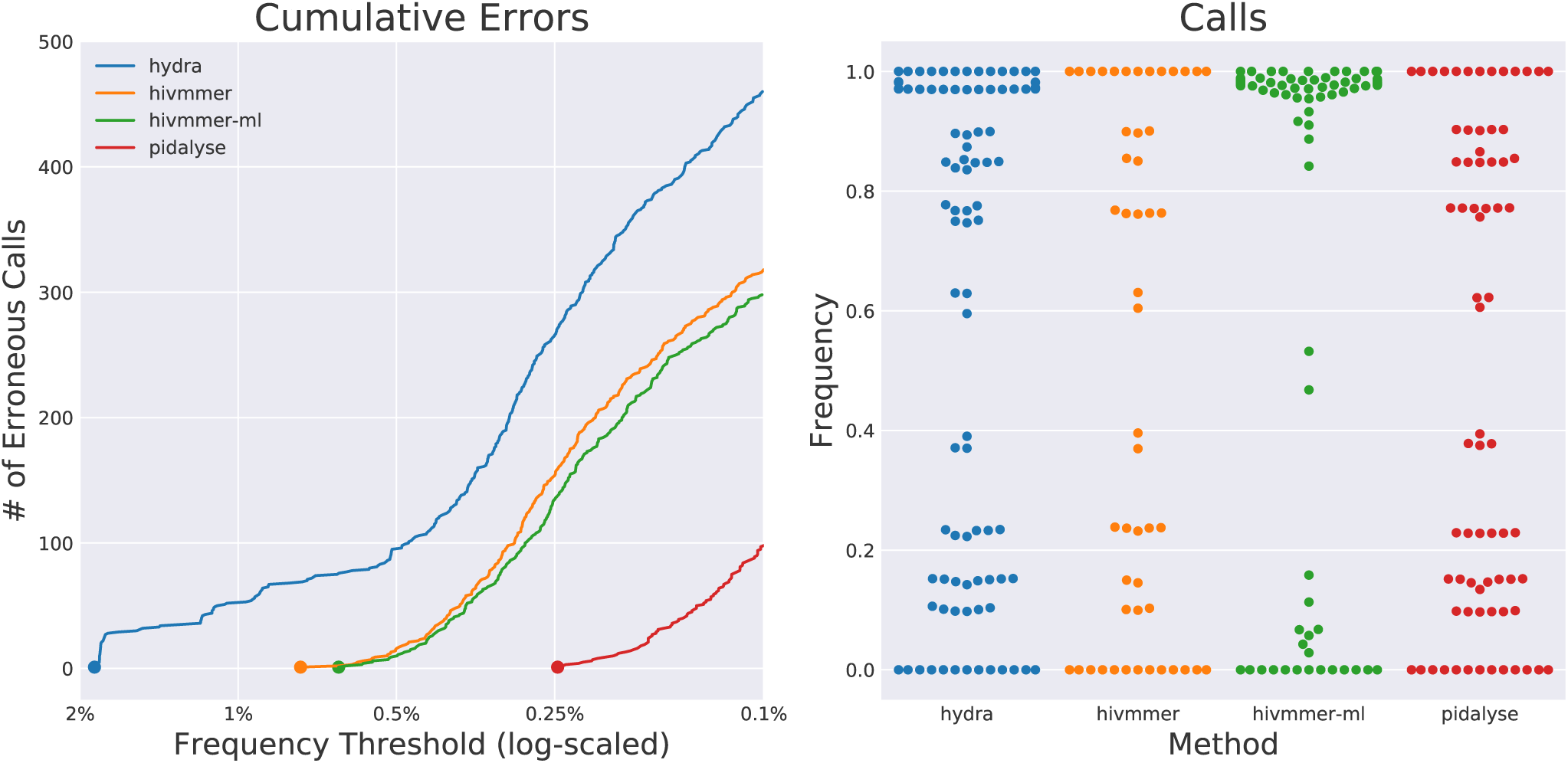
The cumulative error rates and distribution of variant calls in alignments for the PID data set. The pidalyse method exhibits the fewest alignment errors, since it uses the primer IDs to call consensus sequences for each fragment; hivmmer performs closer to pidalyse than HyDRA, and the addition of the machine-learning model (hivmmer-ml) slightly reduces errors, but also skews the distribution of variant calls relative to the other methods.

We assessed the performance of our machine learning models for predicting erroneous calls in hivmmer’s final alignment. In our cross-validation test, the models trained on PL1:1 and PL1:9 performed exceptionally well. Figure 5 shows the receiver-operating characteristic (ROC) curve for false positive versus true positive rates. The area under the ROC (or AUC) is often used as a metric for model fit, and our scores of *>* 0.9 indicate strong predictive power for the models trained and validated across the PL1:1 and PL1:9 data sets.

**Figure 5.**
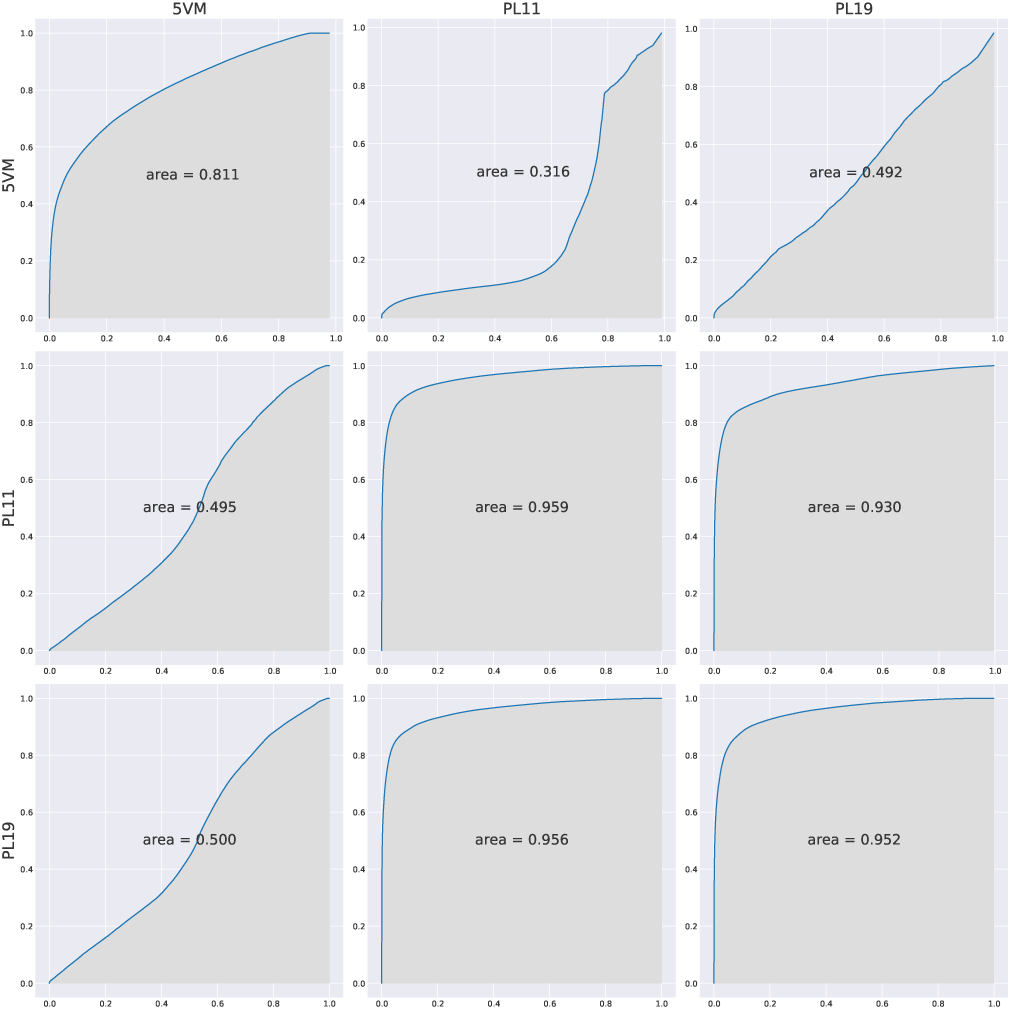
Receiver-operating characteristic (ROC) curves and area-under-the-curve (AUC) showing the trade-off between false-positive and true-positive rate for each of the xgboost models, with the training data set specified on the right side and the validation data set on the top. The models cross-validate well between the PL1:1 and PL1:9 data sets, but not between 5VM and the PL data sets.

Finally, we applied the model trained on 5VM to the hivmmer alignments for the PID data set and used it to mask all alignments with a predicted probably of error ¿0.01. We labeled this method “hivmmer-ml” in Figure 4a, and it did perform slightly better than hivmmer alone, although not as well as pidalyse. However, the hivmmer-ml approach also appeared to bias the distribution of calls, as seen in Figure 4b.

## 4 Discussion

We have introduced a new variant-calling pipeline, hivmmer, whose alignments exhibit lower error rates than existing pipelines on deep Illumina sequencing of HIV plasmid data. We also demonstrated how machine-learning classification of errors is a promising future extension of this pipeline.

### 4.1 Global thresholding

Our results validate that in some cases the widely accepted 1% global thresholding method will work as expected, as measured on plasmid data sets and assuming the variant calling pipeline has similar accuracy to the pipelines tested here.

Some studies have conducted their own validation of a global threshold. For example, one of the earliest studies to use the global thresholding approach with Illumina MiSeq data was conducted by Dudley *et al.* (2014), who analyzed HXB2 plasmid sequences to determine a higher threshold of 2%. Fisher *et al.* (2015) conducted additional validation for the occurrence of NNRTI drug-resistance mutations at frequencies over 1% in their study using 250 clonal sequences. They also presented a method for error correction using a Bayesian Dirichlet mixture of multinomials probabilistic model to distinguish sequencing error from true low-frequency variants at posterior probabilities *≥*99.99%.

Ode *et al.* (2015) developed an adaptive threshold approach for Illumina MiSeq sequencing based on per-site quality scores and demonstrated that it could reduce mismatches to below a frequency of 1% at most sites on varying mixtures of pNL4-3 and pNL101 plasmid sequences. Their method computes an average quality score across all reads at a reference site and they threshold the variant calls at that site with average score *≥*20 and frequency *≥*1%.

Others, however, have applied thresholding without validation. Studies by Ekici *et al.* (2014), Pessôa *et al.* (2014), and Pessôa *et al.* (2016) applied thresholds of 1% without providing any citation or methodological justification for this approach to error correction. A review of clinical applications of deep HIV sequencing by Casadellá and Paredes (2016) proposed a rule of thumb of a 1% threshold without citation. However, they emphasized that the frequency threshold is likely limited by the number of RNA molecules in the sample, and caution that the frequencies that are retained between 1-100% are likely skewed by many sources of bias during sequencing, and cannot be treated as a linear scale, which is confirmed in our results where we do not recover the expected frequencies given the plasmid mixture proportions.

In our results, we did find erroneous calls above 1% (HyDRA, 5VM), which call into question the rule of thumb of 1%. Moreover, Ode *et al.* (2015) claimed in their analysis that they found mismatches occurring at as high as 6.4% frequency at some sites. Both their results and ours clearly establish the heterogeneity in error profiles, and that a global threshold is overly conservative at most sites.

Going forward, studies using deep Illumina sequencing of HIV to analyze variants at low frequencies should include control data sets and detailed analysis of the error profile, such as the one we have presented in this study. One potential study design is to multiplex a plasmid mixture control into every lane; the control can then be used to establish an error profile for the other HIV samples of interest in that lane.

## 4.2 Machine learning

Global thresholding is, effectively, a crude model that assumes homogenous error profiles at all sites. Ode *et al.* (2015) overcame this limitation by applying an adaptive threshold and modeling the quality scores of the reads. Still, an even more promising approach to modeling the heterogeneity of error profiles is to use a machine learning algorithm that can learn the signature of erroneous calls using more than just the quality scores.

An early example of this approach was given by Meacham *et al.* (2011), who used Illumina GAIIx data from a human genome methyl-Seq experiment to train a logistic regression for classifying errors, specifically focused on GGT motifs. Like in HIV deep sequencing, methyl-Seq also tends to generate many short fragments that yield overlapping reads. Using an alignment to a reference genome, they classified two types of errors: the case where one of the overlapping reads disagrees with the reference and the case where both of the reads disagree. The variables for the regression included the directionality of the read, the quality scores, and the position of the base call in the read.

In the context of HIV, Li *et al.* (2014) performed variantcalling for the *int* gene after removing sequencing errors using a random-forest classifier that they commissioned through a crowd-sourced TopCoder competition. The top entry in the competition used variables computed from the quality scores, alignment scores, and 13-mer frequency of single-end Illumina HiSeq data. They estimated the limit of detection of 0.095%, after applying a threshold on the classifier prediction that removed 99% of false-positive variants in a control library of mixed plasmid sequences at concentrations as low as 0.1%. Our approach extends this by cross-validating a similar model across multiple plasmid data sets. An important finding is that while the models cross-validate well between the PL1:1 and PL1:9 data sets, the cross-validation between those data sets and 5VM was poor, suggesting over-fitting. PL1:1 and PL1:9 contain the same plasmids, were prepared by the same lab, and sequenced within the same lane of an Illumina MiSeq run. Therefore, it appears that predictive modeling of sequencing errors can be applied successfully within a lane, suggesting that future HIV-1 studies may benefit from multiplexing a plasmid mixture into each lane of sequencing.

## 4.3 Overlapping reads as technical replicates

One potential reason why hivmmer outperforms the other pipelines is that it more closely models the fragment distribution through its use of the PEAR read merger. As shown in Figure 1, the majority of reads are complete overlapping (e.g. technical replicates) or partially overlapping in deep Illumina sequencing of HIV. In particular, the fragment distribution is non-normal, while many short-read aligners, including bowtie2, assume a normal distribution of fragments.

PEAR is able to use this replicate information to correct errors at sites where the replicates disagree, by comparing quality scores. This approach has not, to our knowledge, been applied to HIV before, although it was tested by Chen-Harris *et al.* (2013) in a study with 1kb regions of the rabies and BCV viruses. They showed that the PCR error rate exceeds the sequencing error rate at high enough quality scores, and they called variants using a position-dependent model to determine an optimal quality score threshold.

Preston *et al.* (2016) developed a similar protocol called Paired-End Low Error Sequencing (PELE-Seq) that combines barcoding with overlapping read pairs to correct for both PCR and sequencing error and accurately detect rare variants. Although that specific protocol has only been tested with E. coli and nematode DNA samples, the concept is directly relevant to HIV, where barcoding is already in use through the Primer ID protocol.

## 4.4 Primer ID

Our results confirm that the consensus sequences generated by the Primer ID method do achieve lower error rates than any of the pipelines. Primer ID is an area of active research, and most recently Boltz *et al.* (2016) extended the existing methods by using shorter PCR primers and more stringent consensus criteria, in a method they call ultrasensitive single-genome sequencing (uSGS). In comparisons with the earlier methods from Jabara *et al.* (2011), Zhou *et al.* (2015) and Seifert *et al.* (2016), they found that the uSGS technique yielded more unique Primer IDs and overall consensus sequences.

However, an important limitation of all of the Primer ID techniques is the difficulty of multiplexing multiple samples in the same lane, which is a common practice to reduce sequencing cost. In fact, because of the short length of the HIV genome, sufficient depth of coverage can be achieved with many fewer reads than a full lane of Illumina sequencing provides. In the extreme case, this was demonstrated with the successful application of “wide” sequencing by Lapointe *et al.* (2015) to sequence a region of the *pol* gene from 1,143 patient samples in a single Illumina MiSeq run.

In situations where the cost of Primer ID is prohibitive, there are still other avenues for controlling RT-PCR error. Orton *et al.* (2015) developed a computational model for the accumulation of errors following multiple PCR cycles. They validated this model using Illumina GAIIx sequencing of FMDV (not HIV) plasmid sequences with varying rounds of PCR amplification, including a condition with no amplification, and found that RT-PCR errors were concentrated in specific areas related to known variability in the FMDV genome, and not evenly distributed across the genome. They also found that most of the error came from the PCR amplification rather than the RT step in sample preparation. Overall, their recommendation is to use the highest fidelity enzymes and minimize the number of PCR cycles.

Zanini *et al.* (2016) presented an Illumina MiSeq protocol with single-round PCR and a new primer design for HIV, and found an error rate of 0.1% that they attribute to PCR error, after removing low quality base calls. They validated the correlation between base calling errors and quality scores with a PhiX spike-in. Furthermore, they tested for in-vitro recombination and found it in nearly 10% of reads generated from nested PCR, but almost none in those from single-round PCR.

Thus, a viable alternative to Primer ID in future experiments may be to combine a sequencing protocol using high fidelity enzymes and single-round PCR with hivmmer.

## 5 Conclusion

The ideal sequencing technology for genomic studies of HIV would generate full-length reads, without error, of individual virus particles from a patient. Although this is not technically possible with today’s technology, understanding the causes and corrections for measurement errors and optimizing ways to avoid them will get us closer to that goal. Newer, longer-read and single-molecule sequencing technologies such as PacBio and Oxford Nanopore also hold promise in addressing these limitations, although they currently have much higher error rates than Illumina sequencing (Goodwin *et al.*, 2016). Thus, even with newer and improved sequencing technology, understanding measurement error will still be a priority for making robust inferences from HIV sequencing data.

However, a concern with all of the approaches to variantcalling presented here, both thresholding and machine-learning, is the potential of overfitting a method to the training and validation data. Any robust method will need to be cross-validated on a wider variety of HIV data before it can be trusted as a general-purpose tool. It will be important to benchmark error correction methods on datasets from different labs, that use varying protocols to sequence varying mixtures of plasmid sequences and regions of the HIV genome. This can be facilitated by more public sharing of plasmid datasets, several of which have already been deposited with the Sequence Read Archive.

Further refinement of error correction methods for deep Illumina sequencing of HIV that combines protocols to reduce PCR errors with overlapping read pairs and machine learning classification of sequencing errors will be a valuable and important step toward more robust, and ultimately clinically-trusted, tools for HIV genotyping. Overall, these clear directions for future work will benefit the HIV research community by enabling more robust inference. In the specific context of drug resistance mutations, more robust error correction will allow for more sensitive detection of emerging resistance at very low variant frequencies and the continued exploration of their significance.

## Acknowledgements

This research was conducted using computational resources and services at the Center for Computation and Visualization, Brown University.

---

**Figure S1.**
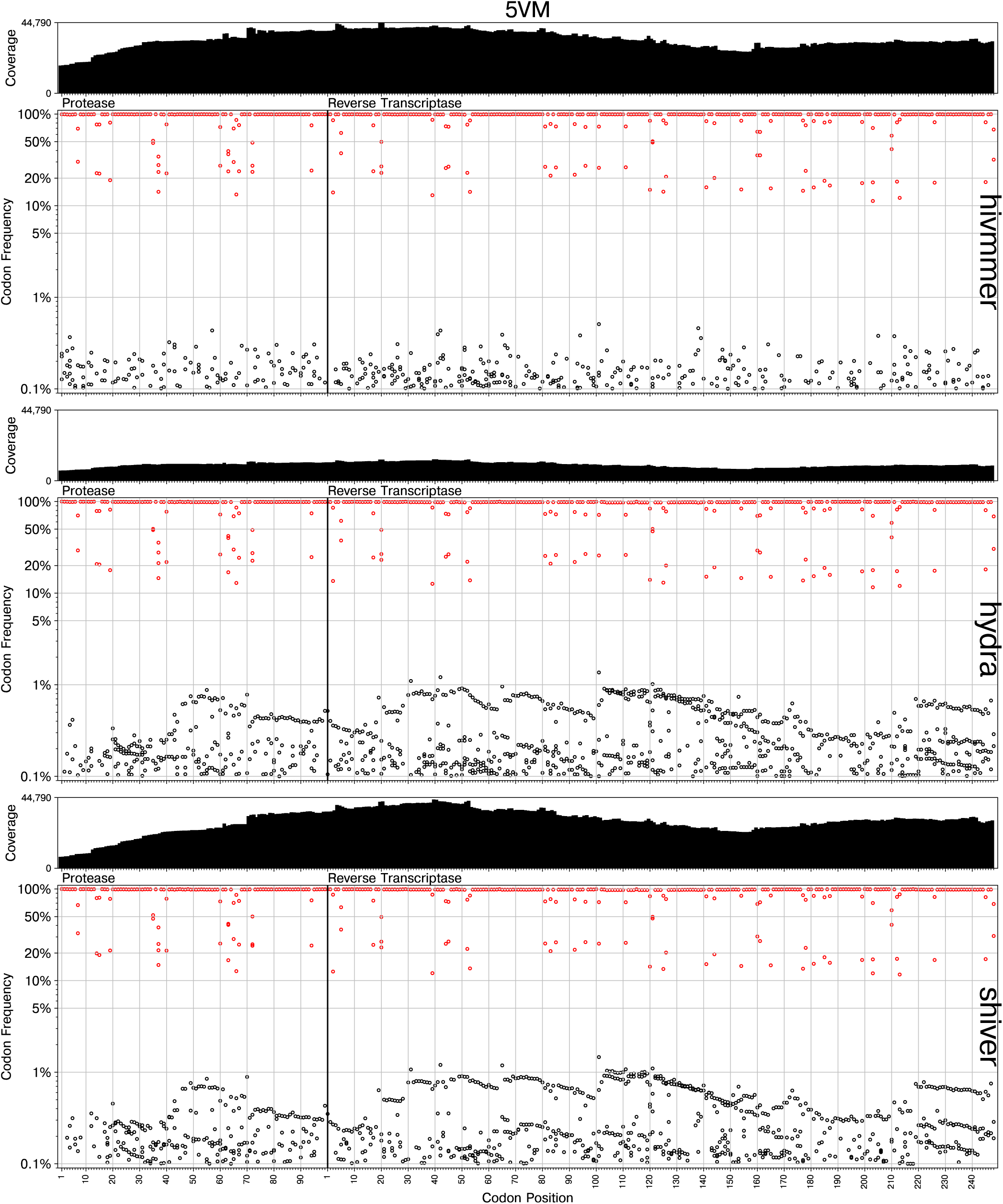
Detailed view of variant calls for the 5VM data set.

**Figure S2.**
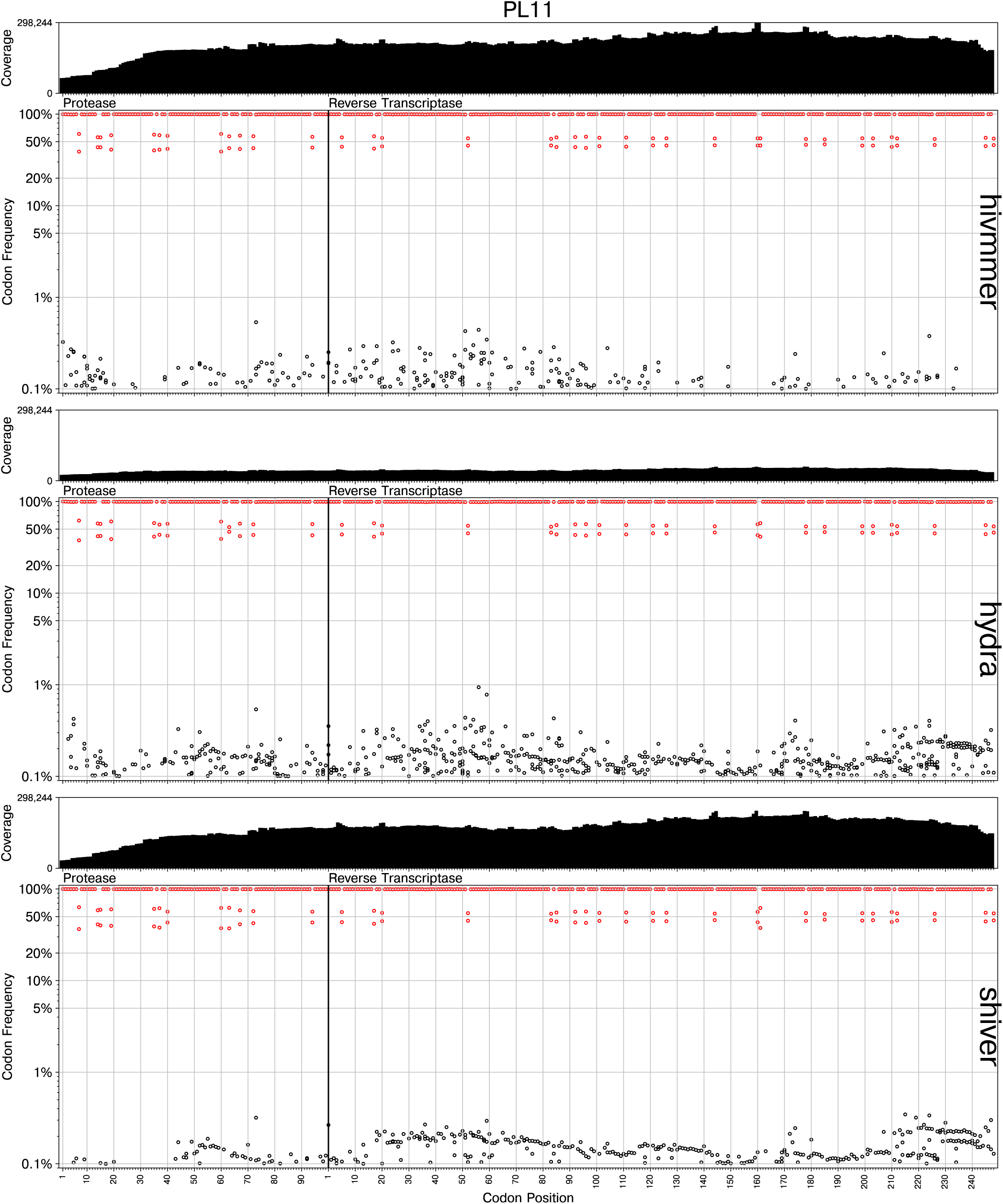
Detailed view of variant calls for the PL1:1 data set.

**Figure S3.**
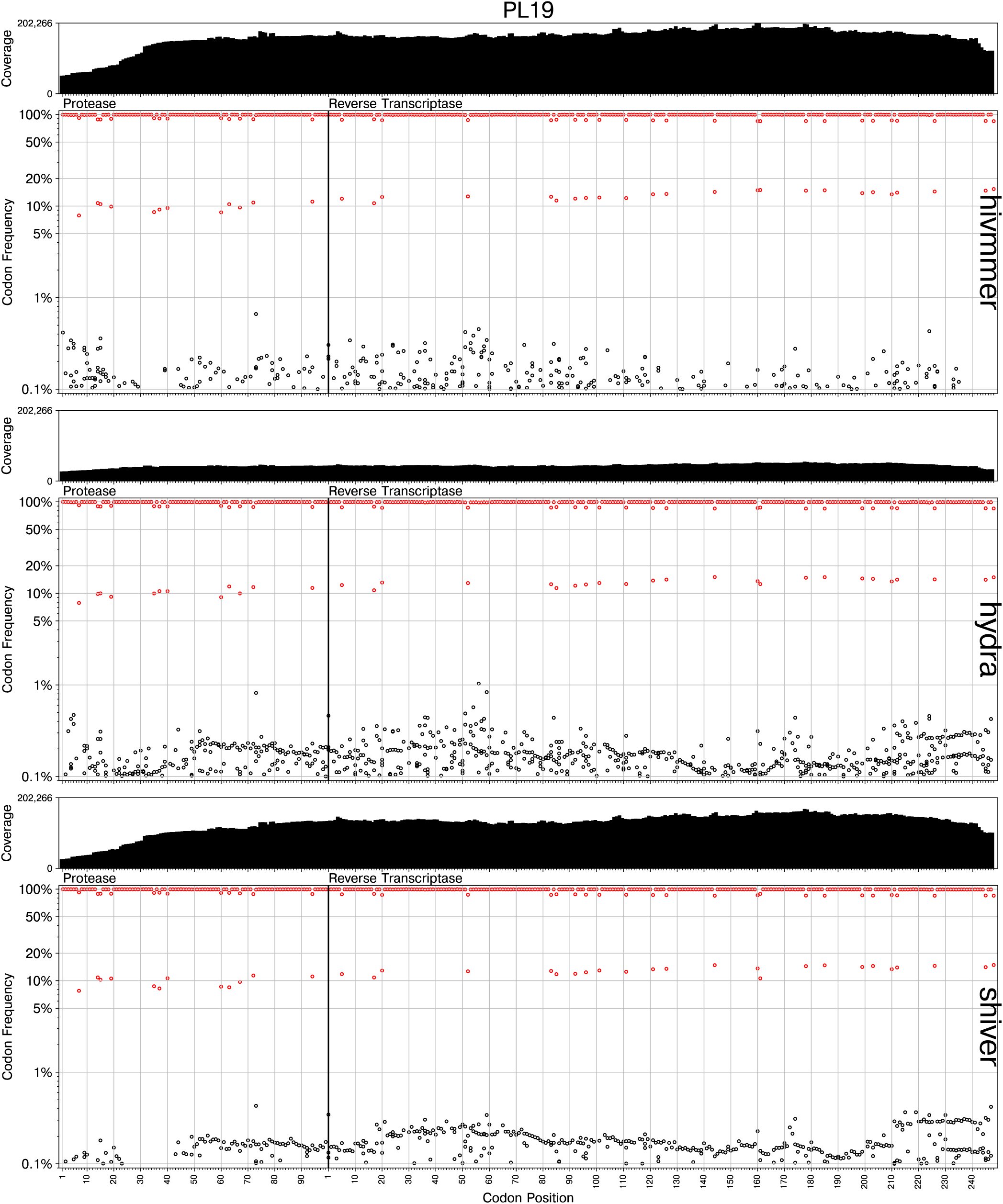
Detailed view of variant calls for the PL1:9 data set.

**Figure S4.**
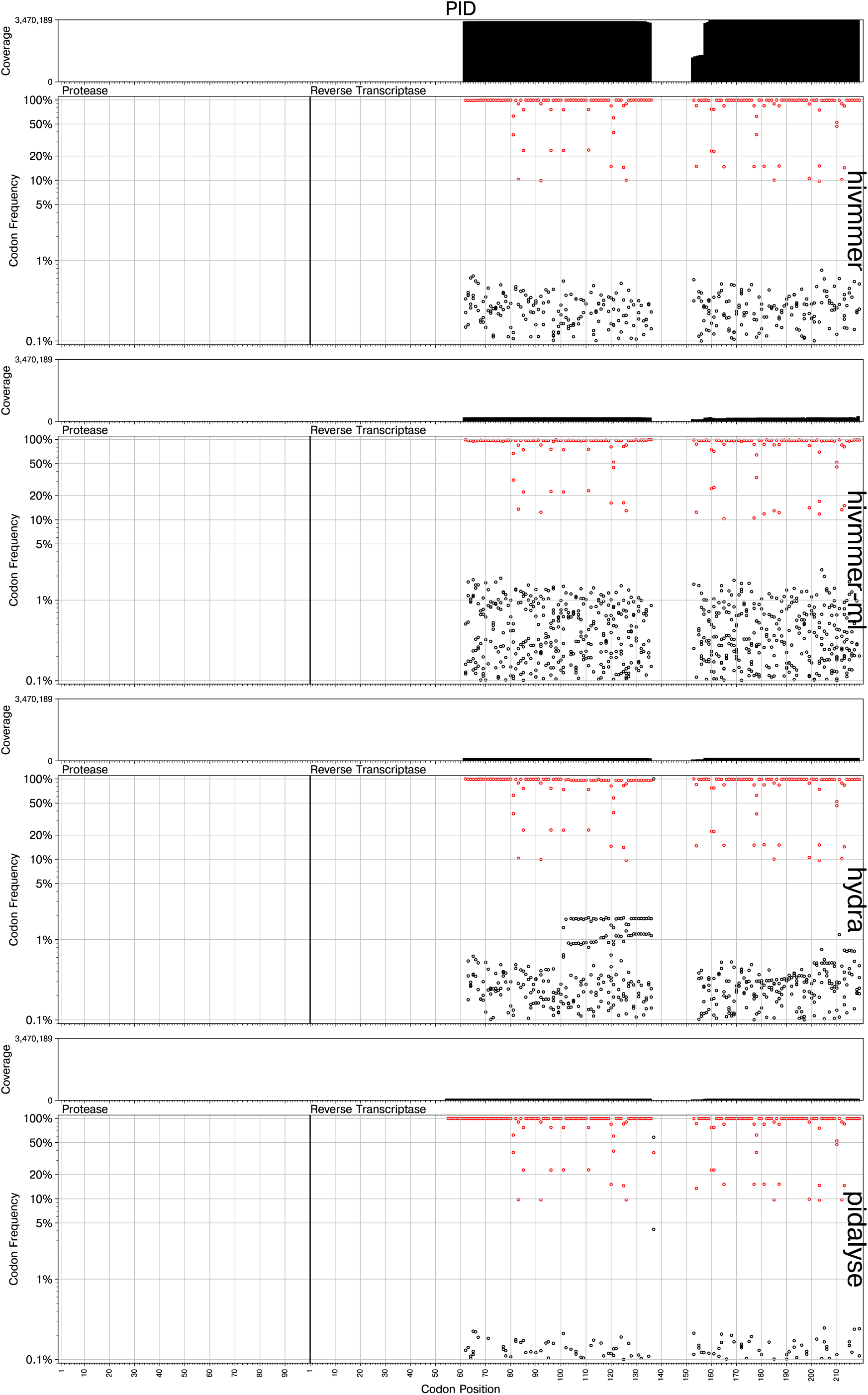
Detailed view of variant calls for the PID data set.

